# Global metabolic reprogramming and sex-specific impact of caloric restriction in a mouse model of Alzheimer’s disease

**DOI:** 10.1101/2025.05.24.655912

**Authors:** Myroslava V. Vatashchuk, Viktoriia V. Hurza, Maria M. Bayliak, Dmytro V. Gospodaryov, Volodymyr I. Lushchak, Olga Garaschuk

**Author notes:** Corresponding authors: Olga Garaschuk Volodymyr I. Lushchak Dmytro V. Gospodaryov Maria M. Bayliak.

## Abstract

**Background:** Alzheimer’s disease (AD), disproportionally affecting women, is generally regarded as a disease of the brain. Yet, cumulative evidence increasingly supports a more systemic, full-body view on this incurable disorder, with the liver and kidneys playing an important role in amyloid clearance. The latter is likely potentiated by caloric restriction (CR), but its impact on the metabolism of key amyloid-handling tissues is poorly understood.

**Methods:** We examined the sex-specific effects of amyloidosis and CR on oxidative and metabolic processes in APPPS1 mice, expressing mutant amyloidogenic proteins under the neuronal Thy-1 promoter. Wild-type (WT) and APPPS1 mice were either fed *ad libitum* (AL) or received 70% of their individual AL intake (CR regimen).

**Results:** Compared to age-matched WT controls, the brain, liver, and kidney of 9-month-old AL APPPS1 mice exhibited higher levels of oxidative stress markers along with altered activity of antioxidant enzymes, including higher superoxide dismutase and lower catalase activity. These differences were sex- and tissue-specific, with kidneys showing the largest AD-induced differences between sexes. As for key glycolytic enzymes, APPPS1 mice possessed higher activity of pyruvate kinase than WT mice in all organs and higher hexokinase and phosphofructokinase activities in the brain, with stronger effects observed in males. CR intensified oxidative stress in the liver and the female brain but reduced it in the female kidney. Moreover, it potentiated glycolysis predominantly in females and modulated glutathione-dependent enzymes in a sex-dependent manner.

**Conclusions:** Neuron-driven amyloidosis induces strong and significant metabolic changes in the entire body, encompassing ubiquitous oxidative damage, enhancement of glycolysis, and antioxidant defense. The lower activity of glycolytic enzymes in the female brain and decreased antioxidant defense in the female liver and kidney provide yet another facet to the complex picture of female vulnerability in AD. Females seemed to profit more from CR, but the general CR effect was not very strong.

## 1. Background

Alzheimer’s disease (AD) is the most widespread neurodegenerative disorder, characterized by progressive memory loss, cognitive decline, and changes in behavior and personality. A predominant sporadic form of the disease (late-onset AD, LOAD) is likely caused by a combination of genetic, environmental, and lifestyle factors and typically develops after the age of 65. The early-onset AD comprises about 2-10% of all AD cases and is typically associated with autosomal-dominant mutations in genes encoding amyloid precursor protein (APP) or presenilins 1 (PS1) and 2 (PS2) [1,2]. Nearly two-thirds of individuals diagnosed with AD are women, with AD ranking as the fifth leading cause of death for women (6.1% of deaths) and seventh (2.6% of deaths) for men [3].

Histologically, AD is defined by the presence of extracellular amyloid deposits (senile plaques built of amyloid β (Aβ)) in the brain parenchyma and intracellular accumulations of neurofibrillary tangles containing hyperphosphorylated microtubule-associated protein tau. The functional hallmarks of the disease include synaptic vulnerability, metabolic dyshomeostasis, impaired neuro-glial Ca^2+^ signaling, endosomal/lysosomal dysfunction, impairment of autophagy, activation of the brain’s innate immune system, oxidative stress/damage, as well as impaired lipid metabolism [2,4–6].

Interestingly, APP expression is not restricted to the brain but is also found in the peripheral tissues like skin, intestine, and liver [7]. The same applies to amyloid plaques, which, for example, have been detected in the gastrointestinal tracts of AD patients and mouse models of AD. Moreover, peripherally generated Aβ is capable of entering the brain and aggregating therein, as documented by a parabiosis model between AD (APP/PS1) and WT mice or a mouse model selectively overexpressing human APP with Swedish (KM670/671NL) and Indiana (V717F) AD mutations in hepatocytes [8,9]. Besides, peripheral organs such as the liver, kidney, spleen, and gut are important for Aβ clearance [7,10].

The progression of AD is accompanied by the impairment of energy metabolism [5]. In fact, inefficient glucose utilization represents one of the early and well-known hallmarks of AD, which often comes hand-in-hand with peripheral (and likely also central) insulin resistance [11]. One possible mechanism promoting insulin resistance is the AD-driven hyperactivation of the mechanistic target of rapamycin (mTOR). Activation of the mTOR pathway is known to impact protein biosynthesis, mitochondrial function, lipo- and ketogenesis, and to inhibit autophagy and insulin signaling [5]. It is crucially linked to glucose metabolism and was shown to increase the production of reactive oxygen species (ROS), tau hyperphosphorylation, cerebrovascular dysfunction, and likely also neuroinflammation [12]. Sirtuin-3 (SIRT3), a mitochondrial nicotinamide adenine dinucleotide (NAD^+^)-dependent protein deacetylase, is another key player impacting mitochondrial function, lipid metabolism, ROS production, inflammation, and antioxidant defense. SIRT3 is widely expressed in metabolically active tissues, including the heart, liver, kidney, and brain, and SIRT3 expression is downregulated in the brain tissue of AD patients and mouse models of AD [13]. Cumulative evidence suggests that increasing SIRT3 activity might alleviate AD progression by positively influencing mitochondrial homeostasis and neuroinflammation, activating antioxidant enzymes, and decreasing levels of tau and Aβ.

Several studies have shown that calorie restriction can attenuate the severity of AD symptoms by reducing the density of amyloid plaques and inhibiting mTOR signaling, generation of ROS, neuroinflammation and tau hyperphosphorylation as well as increasing SIRT3 expression in the liver and brain [14–19]. Yet, the opposite effects were also reported [20–22]. For example, the every-other-day feeding of 5XFAD transgenic female mice during the plaque deposition period (2-6 months of age) failed to decrease plaque size or density, brain Aβ42 levels, and the AD-induced permeability of the blood-brain barrier [21]. Moreover, it exacerbated the activation of microglia and astrocytes, increased the brain level of proinflammatory cytokine tumor necrosis factor α, further reduced the density of synaptic marker synaptophysin, and promoted neuronal death. Similarly, no effect of caloric restriction (CR) on amyloid plaque density was found in the aged rhesus monkey population fed 30% less than individual sex-, age-, and weight-matched controls for up to 15 years (till death [22]), in line with the relatively low WT mouse brain responsiveness to CR [23–26].

In the current study, we explored the amyloidosis-induced response of key amyloid-handling organs (brain, liver, and kidney) in a widely used mouse model of AD (APPPS1 mice [27]) and tested the ability of CR to mitigate oxidative stress caused by the accumulation of Aβ. In calorie-restricted and *ad libitum*-fed APPPS1 (hereafter AD) mice as well as in age-matched WT littermates, we measured markers of oxidative stress and the activities of antioxidant enzymes, as well as activities of key glycolytic enzymes and lactate dehydrogenase (Fig. 1A) to infer metabolic changes caused by both amyloid accumulation and CR. The analyses considered sex as a biological variable, thus deepening our understanding of possible mechanisms underlying the increased incidence rates for Alzheimer’s disease in women [28].

**Figure 1.**
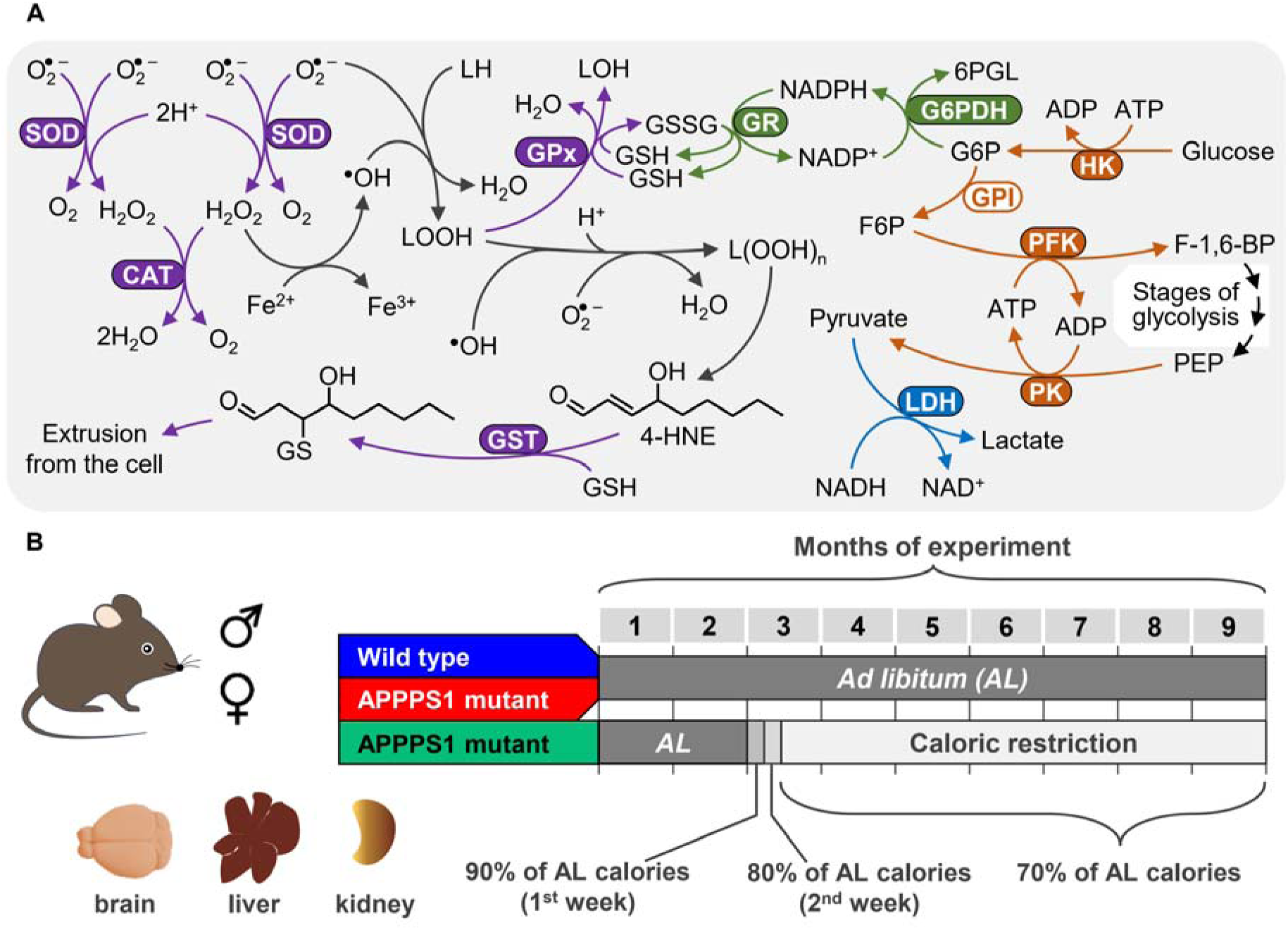
Experimental settings. A, Scheme illustrating the relationship between oxidative damage, antioxidant defense enzymes, and glycolytic enzymes. Reactive oxygen species (ROS), such as superoxide anion radical (O_2_^•^[) and hydrogen peroxide (H_2_O_2_), are metabolized by superoxide dismutase (SOD), catalase (CAT), and glutathione peroxidase (GPx), with the support of glutathione reductase (GR) and NADPH. The latter is produced in glucose-6-phosphate dehydrogenase (G6PDH) reaction during the oxidation of glucose-6-phosphate (G6P) to 6-phosphoglucolactone (6PGL). G6P is formed in the reaction of glucose phosphorylation catalyzed by hexokinase (HK) and can enter glycolysis. Hexokinase (HK), and phosphofructokinase (PFK) and pyruvate kinase are key enzymes of glycolysis, which catalyze irreversible reactions. Lactate dehydrogenase (LDH) converts pyruvate to lactate to maintain redox balance. If the activity of antioxidant enzymes and the pool of reduced glutathione (GSH) are insufficient, oxidative damage to proteins and lipids increases. The scheme shows how ROS initiate lipid peroxidation and emphasizes the importance of GSH in forming primary and relatively stable products of lipid oxidation, lipid hydroperoxides (LOOH). Activities of enzymes whose names are shown in white font on the colored background were measured in the present study (all except GPI – glucose-6-phosphate isomerase). Metabolite abbreviations are as follows: F6P – fructose-6-phosphate, F-1,6-BP – fructose-1,6-biphosphate, PEP – phosphoenolpyruvate, 4-HNE – 4-hydroxy-2-nonenal. Arrows and enzymes shown in grape color denote reactions directed to antioxidant defense; grey arrows mark reactions that promote oxidative damage in the form of lipid peroxides and products of their breakdown. Enzyme (GST) and arrows shown in liver color mark antioxidant defense-related conjugation and excretion of 4HNE. Green arrows and enzymes mark reactions directed to glutathione reduction, tawny-colored enzymes and arrows – key glycolytic reactions, and cyan arrow – the lactate dehydrogenase reaction. B, Experimental arrangement as detailed in Materials and methods.

## 2. Materials and methods

### 2.1 Animals and experimental conditions

C57BL/6N (wild type, WT) and APPPS1 mice of both sexes were used. APPPS1 mice overexpress human APP with the Swedish mutation (KM670/671NL) and presenilin 1 with the L166P mutation under control of the neuronal Thy-1 promoter. In these mice, both the PS1 and the APP constructs are restricted in expression to the postnatal brain, and cerebral amyloidosis starts at 6-8 weeks of age [27].

WT mice (3 males and 4 females) and one group of APPPS1 mice (AL group, 4 males and 5 females) had unlimited access to food and water during 9 months of the experiment (Fig. 1B). The other group of APPPS1 mice (CR group, 4 males and 4 females) had unrestricted access to food for two months and thereafter was switched to caloric restriction for seven months, until the end of the experiment. In the CR group, each mouse received 90% of its individual *ad libitum* food intake during the first week and 80% during the second week of the CR regimen. Starting from the third week and continuing until the end of the experiment, the CR animals received 70% of their respective *ad libitum* food intake. All animals were kept under standard conditions with a 12 h light/dark cycle. *Ad libitum*-fed mice stayed in groups of 3–5 mice, and CR mice were kept individually. All procedures complied with the ARRIVE guidelines, were carried out following the EU Directive 2010/63/EU for animal experiments, and were approved by the state government of Baden-Württemberg, Germany.

### 2.2 Tissue sampling

Mice were deeply anesthetized (ketamine 200 mg/kg body mass; xylazine 20 mg/kg body mass), tested for the surgical depth of anesthesia by checking the interphalangeal reflex, and subjected to the transcardiac perfusion with cold phosphate-buffered saline (PBS) until blood vessels and organs were largely bloodless [29]. After collection, the liver, kidneys, and brain (without olfactory bulb and cerebellum) were immediately frozen on dry ice and stored in a freezer at −80 °C.

### 2.3 Tissue homogenization and determination of oxidative stress markers

Lipid peroxides and protein carbonyls were measured as markers of oxidative stress. Lipid peroxides (LOOH) from frozen tissues were extracted with ethanol as described previously [26] and measured with Fe^2+^/xylenol orange method [30,31]. Cumene hydroperoxide was used to build the standard calibration curve [30,31].

To determine the levels of carbonyl groups in proteins, frozen tissue samples were homogenized in 50 mM potassium phosphate buffer (KPi, pH 7.0), containing 0.5 mM *N*,*N*,*N*′,*N*′-ethylenediaminetetraacetic acid (EDTA), and 1 mM phenylmethylsulfonyl fluoride at a ratio of 1:10 (milligram of tissue: microliter of homogenization medium). The homogenates were centrifuged at 21000×*g* for 15 min at 4 °C in a Thermo Fisher Scientific Multifuge X1R centrifuge (Waltham, USA). The supernatants were then mixed with 20% (final concentration) trichloroacetic acid to precipitate proteins. The content of carbonyl groups in proteins was determined by their reaction with 2,4-dinitrophenylhydrazine, resulting in the formation of colored dinitrophenylhydrazones and recorded at 370 nm [32,33].

### 2.4. Tissue homogenization and determination of the activities of antioxidant enzymes

Supernatants were prepared as described above for the carbonyl protein assay. Superoxide dismutase (SOD) activity was measured based on the inhibition of quercetin oxidation by superoxide anion radicals, which were generated in an alkaline medium by TEMED (*N*,*N*,*N*′,*N*′-tetramethylethylenediamine) in the presence of oxygen. The rate of the quercetin oxidation reaction was monitored spectrophotometrically at the 406 nm wavelength [34]. The reaction was run in a mixture containing 30 mM Tris-HCl buffer (pH 10.0), 0.5 mM EDTA, 0.8 mM TEMED, 0.05 mM quercetin, and 0.3-100 μl supernatant; the change in absorbance with time was tested for 6–8 different volumes of supernatant. One unit of SOD activity was defined as the amount of enzyme (per mg protein) that inhibits the quercetin oxidation reaction by 50% of the maximum. Calculations were performed using KINETICS software [35].

The activities of catalase, glutathione peroxidase (GPx), glutathione reductase (GR), glutathione S-transferase (GST), and glucose-6-phosphate dehydrogenase (G6PDH)) were measured spectrophotometrically as described in our previous studies [23,33]. Briefly, all reaction mixtures contained 50 mM KPi (pH 7.0) and 0.5 mM EDTA, and additional components as follows: for catalase – 10 mM H_2_O_2_*, and 10 μl supernatant; for GPx – 0.25 mM NADPH, 4 mM sodium azide, 1 unit (U) GR, 15 mM reduced glutathione (GSH), 0.2 mM H_2_O_2_, and 20 μl supernatant*; for GST – 5 mM GSH, 1 mM 1-chloro-2,4-dinitrobenzene, and 2–5 μl supernatant*; for GR – 0.25 mM NADPH, 1 mM oxidized glutathione (GSSG)*, and 2–50 μl supernatant; for G6PDH – 5 mM MgCl_2_, 0.2 mM NADP^+^, 2 mM G6P* with 20 μl supernatant. Catalase activity was measured by following the rate of H_2_O_2_ decomposition at 240 nm (molar extinction coefficient ε = 39.4 M^−1^ cm^−1^), GST activity was monitored at 340 nm by the formation of 1-(*S*-glutathionyl)-2,4-dinitrobenzene (ε = 9600 M^−1^ cm^−1^), an adduct between GSH and 1-chloro-2,4-dinitrobenzene. Activities of GPx, GR, and G6PDH were assayed at 340 nm by monitoring NADPH (ε = 6220 M^−1^ cm^−1^), consumed (GPx, GR), or formed (G6PDH) in these reactions. Asterisks mark components that were omitted in blank reactions. One unit of catalase, GPx, GST, GR, and G6PDH activity is defined as the amount of the enzyme consuming 1 μmol substrate or generating 1 μmol product per minute.

### 2.5 Tissue homogenization and determination of the activity of glycolytic enzymes

To determine the activity of glycolytic enzymes, frozen tissue samples were homogenized in a lysis medium containing 50 mM imidazole buffer (pH 7.5), 0.5 mM EDTA, 1 mM phenylmethylsulfonyl fluoride, 1 mM dithiothreitol, 20 mM NaF, and 150 mM KCl. The resulting homogenates were centrifuged at 21000×*g* for 15 min at 4 °C. After centrifugation, the supernatants were collected and stored on ice for several hours to determine biochemical parameters.

The activities of hexokinase (HK), phosphofructokinase (PFK), and pyruvate kinase (PK) were measured spectrophotometrically in coupled enzyme reactions by registering NADP^+^ reduction (for HK) or NADH oxidation (for PFK and PK) at 340 nm [25,36]. All assays except LDH activity were run in 50 mM imidazole buffer (pH 7.5) at 25°C with a final volume of 1.0 ml, including supernatant. Blank activities were assayed in mixtures without a specific substrate indicated by the asterisk. LDH activity was measured in 50 mM KPi (pH 7.5). Briefly, reaction buffers contained as follows: for HK – 10 mM glucose*, 0.2 mM NADP^+^, 2 mM ATP, 5 mM MgCl_2_, 0.5 units (U) G6PDH, and 30 μl supernatant, for PFK – 5 mM fructose 6-phosphate*, 5 mM MgCl_2_, 5 mM ATP, 0.16 mM NADH, 50 mM KCl, 0.5 U aldolase, 0.5 U triose-phosphate isomerase, 2 U glyceraldehyde-3-phosphate dehydrogenase, and 30 μl supernatant, for PK – 1 mM 2-phosphoenolpyruvate*, 5 mM MgCl_2_, 50 mM KCl, 2.5 mM ADP, 0.16 mM NADH, 2.5 U LDH, and 5 μl supernatant. The reaction mixture for lactate dehydrogenase contained 0.5 mM EDTA, 0.2 mM NADH, 1 mM pyruvate*, and 10 μl supernatant. The total protein concentration in the samples was determined by the Bradford method [37] with bovine serum albumin as a standard.

### 2.6 Statistical analysis

The normality of the datasets was tested using the Jarque-Bera test and the homoscedasticity was tested using the Brown-Forsythe test, implemented in the R package ‘DescTools’. If necessary, the data were logarithmically transformed. Data were analyzed using generalized linear models (Gaussian distribution and identity link function), followed by the analysis of deviance and multiple comparisons using estimated marginal means. The analyses were conducted in R using packages ‘car’ (for analysis of deviance) and ‘emmeans’ (for multiple comparisons). A combined factor, mutation plus diet, was chosen as a predictor variable. The sex of animals was chosen as another factor. Interaction between the combined factor and sex was included in the model.

Data are represented as box-and-whisker plots with marked median, first and third quartiles (box floor and ceiling, respectively). The whiskers are extended to the most extreme data points, which do not exceed the 1.5 interquartile range, and other data points are shown as outliers. Differences between the groups for which the *p*-value < 0.05 were considered statistically significant. The boxplots were made using GraphPad Prism version 8.0.2 for Windows (GraphPad Software, Boston, Massachusetts, USA).

Principal component analysis (PCA) was conducted and visualized using R packages ‘ggplot2’, ‘ggpubr’, ‘ggfortify’, ‘ggrepel’, and ‘gridExtra’. The principal components were estimated using an algorithm of a singular value decomposition implemented in the ‘base’ function *prcomp*.

## 3 Results

### 3.1 Lipid peroxide levels and activity of antioxidant and glycolytic enzymes in the brain

In the brains of male and female *ad libitum*-fed AD mice (AL group), the levels of LOOH, a marker of lipid peroxidation intensity, were 3.4- and 2.2-fold (here and below, the median values per group are reported) higher compared to those measured in WT littermates, whereas no difference in lipid peroxides was observed between AL and CR groups of AD mice (Fig. 2A). SOD activity was 2.8- and 2.5-fold higher in male and 3.1- and 3.9-fold higher in female AD mice on AL and CR regimens compared to the values measured in the respective WT groups (Fig. 2B). Under AL regimen, catalase activity was by 56% lower in female AD mice, compared to respective WT mice; with no difference observed between AL and CR groups of AD mice (Fig. 2C).

**Figure 2.**
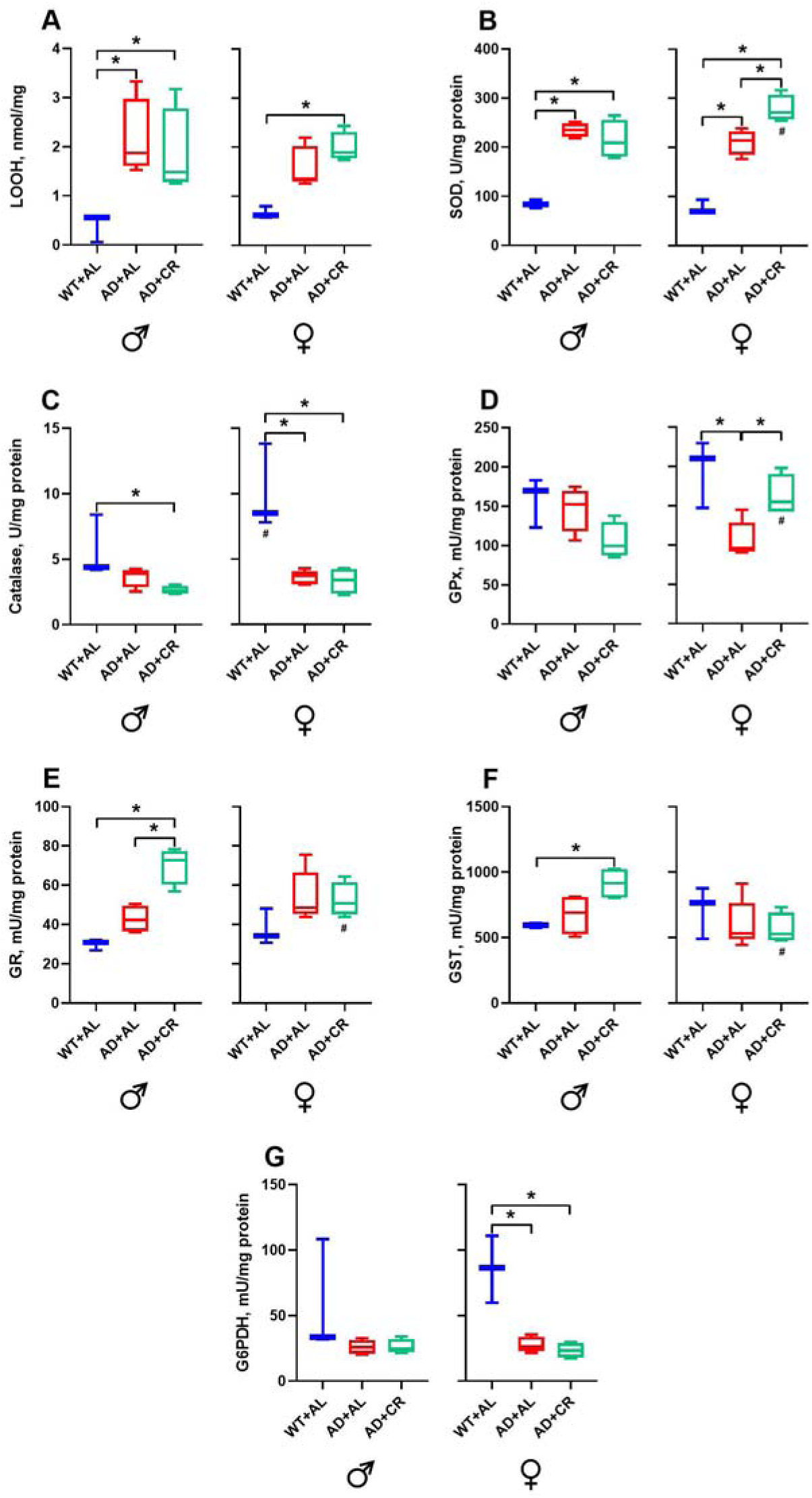
Markers of oxidative stress and activities of antioxidant enzymes in the brain. Effects of amyloidosis (AD phenotype) and caloric restriction (CR) on the APPPS1 (AD) and C57BL/6N (WT) mice of both sexes. Here and below: AL – mice fed *ad libitum*. A –levels of lipid peroxides, B – superoxide dismutase activity, C – catalase activity, D – glutathione peroxidase activity, E – glutathione reductase activity, F – glutathione S-transferase activity, and G – glucose-6-phosphate dehydrogenase activity. Here and below n = 7 WT mice (3 males and 4 females), 9 AL-fed APPPS1 mice (4 males and 5 females) and 8 CR APPPS1 mice (4 males and 4 females). *Significantly different between genotype+diet groups, *p <* 0.05. ^#^Significantly different among sexes, *p* < 0.05.

Male WT and AD mice fed *ad libitum* did not differ in the activities of GPx, GR, GST, and G6PDH (Fig. 2D-G). CR regimen, however, resulted in 2.4-fold higher GR (Fig. 2E) and 1.5-fold higher GST (Fig. 2F) activities in AD males compared to the respective WT groups. In contrast, the GPx (Fig. 2D) and G6PDH (Fig. 2G) activities were significantly lower in the brains of AD compared to WT females, while the GR (Fig. 2E) and GST (Fig. 2F) activities were similar. Under the caloric restriction, females of AD mice had 1.3- and 1.6-fold higher activities of SOD and GPx, respectively, compared to males (Fig. 2B and 2D), while the activities of GR and GST were 30% and 42% lower in AD females compared to males (Fig. 2E and 2F). Note that only in females CR was able to revert the AD-mediated decrease in the GPx activity (Fig. 2D).

*Ad libitum*-fed males of AD mice had significantly (3.4-, 6.3-, 34- and 1.5-fold, respectively) higher HK (Fig. 3A), PFK (Fig. 3B), PK (Fig. 3C), and LDH (Fig. 3D) activities compared to the respective WT male group. In AD males, caloric restriction did not affect HK (Fig. 3A), PFK (Fig. 3B), and LDH (Fig. 3D) activities. However, it significantly reduced the AD-mediated increase in the PK activity (Fig. 3C). Compared to WT females, AD females in the AL group also exhibited significantly (4.3-, 6.2-, 60-, and 1.4-fold, respectively) higher activities of HK (Fig. 3A), PFK (Fig. 3B), PK (Fig. 3C), and LDH (Fig. 3D). Caloric restriction further increased the PFK activity in AD females, leading to significantly different level of PFK activity in males and females, with little impact on the activities of other enzymes tested (Figs. 3A-D).

**Figure 3.**
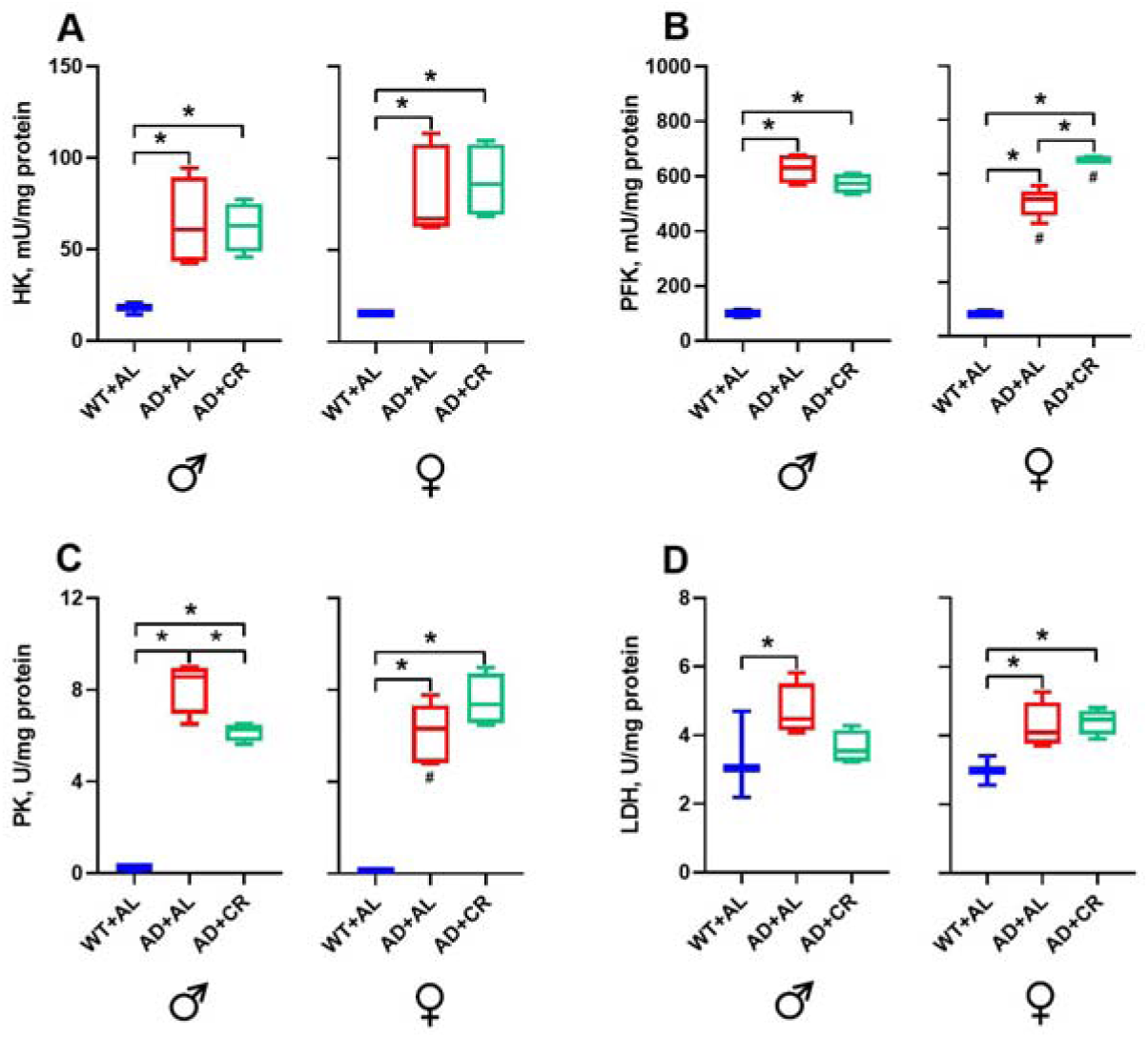
The activities of glycolytic enzymes in the brain. A – hexokinase activity, B – phosphofructokinase activity, C – pyruvate kinase activity, and D – lactate dehydrogenase. *Significantly different between genotype+diet groups, *p <* 0.05. ^#^Significantly different among sexes, *p* < 0.05.

### 3.2 Oxidative stress markers and activities of antioxidant and glycolytic enzymes in the liver

Male AD mice on the AL and CR regimens had 3.7- and 6.7-fold higher levels of LOOH in the liver, compared to the WT group (Fig. 4A). Of note, the levels of lipid peroxides in the liver of AD males increased further (1.8-fold compared to the respective AL group) under caloric restriction (Fig. 4A). Similar situation was observed in AD females. Livers of AL and CR AD females showed 3.7- and 5.2-fold higher LOOH levels than those measured in the WT group. CR group of AD females exhibited 41% higher LOOH levels than the AL counterparts (Fig. 4A). The content of protein carbonyl groups (CP), a marker of protein oxidation, in the male liver was not affected by the amyloidosis or dietary regimen (Fig. 4B). However, AL-fed AD females had by 60% lower CP levels compared to the WT counterparts, likely reflecting the protective effect of Aβ on the liver [38]. SOD activity was significantly (2.2-fold) higher in AL-fed AD compared to WT males (Fig. 4C), and this AD-induced increase was reverted by CR. In contrast, no significant changes in the activity of this enzyme were observed in females. Catalase activity was significantly higher in male compared to female WT mice and decreased in both sexes of AD mice, with the difference becoming significant in male mice only (Fig. 4D).

**Figure 4.**
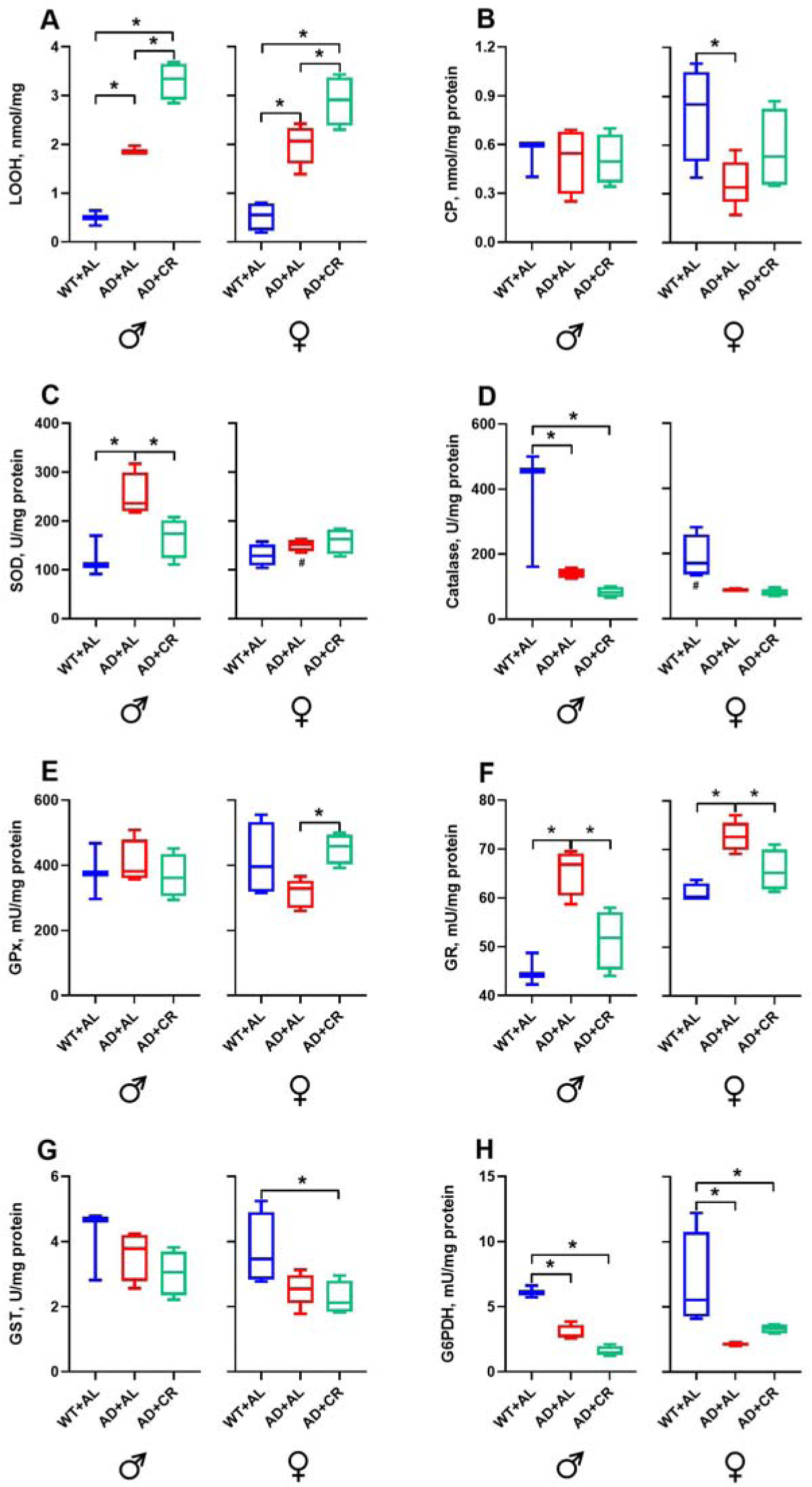
Levels of the oxidative stress markers and activities of antioxidant enzymes in the liver. A – levels of lipid peroxides, B – levels of carbonyl proteins, C – superoxide dismutase activity, D – catalase activity, E – glutathione peroxidase activity, F – glutathione reductase activity, G – glutathione S-transferase activity, and H – glucose-6-phosphate dehydrogenase activity. *Significantly different between genotype+diet groups, *p <* 0.05. ^#^Significantly different among sexes, *p* < 0.05.

AL-fed AD males did not differ from the WT males in GPx (Fig. 4E) and GST (Fig. 4G) activities. The caloric restriction did not affect the activity of these enzymes either. In contrast, GR activity was significantly (1.5-fold) higher in AD and significantly (by 23%) lower in AD+CR male mice (Fig. 4F). Compared to the WT group, liver G6PDH activity was by 55% and 76% lower in AL and CR groups of AD males, respectively (Fig. 4H). Compared to WT littermates, AL*-*fed AD females showed a trend towards reduced activity of GPx and GST and had a significantly (1.6-fold) higher GR as well as a significantly lower (by 62%) G6PDH activity (Fig. 4E-H). Importantly, CR changed the GPx and GR activities significantly, bringing them much closer to control values measured in female WT mice (Fig. 4E, F). No differences between dietary regimens were found for GST and G6PDH activities in female AD mice.

Compared to the respective WT group, AL-fed AD mice of both sexes had significantly (by 4.8-fold in males and 2.6-fold in females) elevated glucose levels, with comparable glucose levels in AL and CR groups of AD mice (Fig. 5A). *Ad libitum*-fed AD males showed similar hepatic HK and LDH activities but a significantly (57%) lower PFK activity and a significantly (2.3-fold) higher PK activity compared to WT males (Fig. 5B-E). AD with or without CR affected neither HK (Fig. 5B) activity in males nor LDH activity (Fig. 5E) in both sexes. In the livers of AL-fed AD females, the activity of HK was slightly (1.3-fold, Fig. 5B) and PK significantly (4.4-fold, Fig. 5D) higher compared to that measured in WT females. Hepatic PFK activity was significantly (43%) lower in the AL-fed AD females compared to the WT group (Fig. 5C). The only difference between dietary regimens was observed for PK, which was significantly higher in CR compared to AL AD females. Moreover, sex-specific differences were observed for HK in the AD+CR group and for LDH in the WT group of mice (Fig. 5B, E).

**Figure 5.**
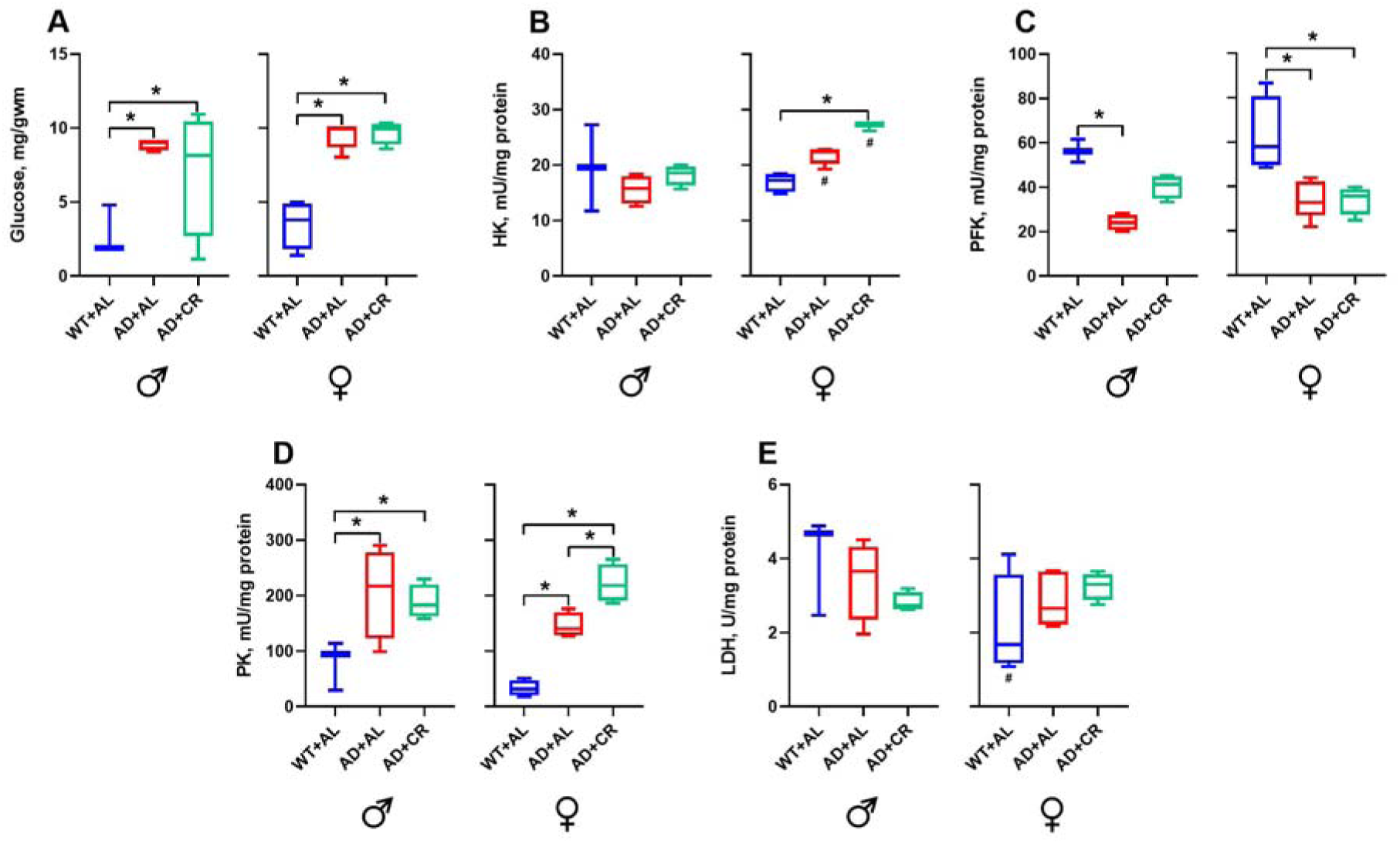
Levels of glucose and activity of glycolytic enzymes in the liver. A – levels of glucose, B – hexokinase activity, C – phosphofructokinase activity, D – pyruvate kinase activity, and E – lactate dehydrogenase activity. *Significantly different between genotype+diet groups, *p <* 0.05. ^#^Significantly different among sexes, *p* < 0.05.

### 3.3 Oxidative stress markers, activities of antioxidant and glycolytic enzymes in the kidneys

Under the AL regimen, kidney LOOH levels in AD mice of both sexes were significantly higher (1.7-fold in males, 2.2-fold in females) than those in the respective WT groups (Fig. 6A). CR regimen brought the LOOH levels closer to the WT values in both sexes causing a slight reduction in the CR compared to AL males and a significant reduction in CR compared to AL females. In AL-fed AD mice, the levels of CP slightly decreased in males and significantly increased in females, resulting in a profound sex-specific difference (Fig. 6B). CR regimen normalized the level of protein carbonyl groups in AD mice of both sexes.

**Figure 6.**
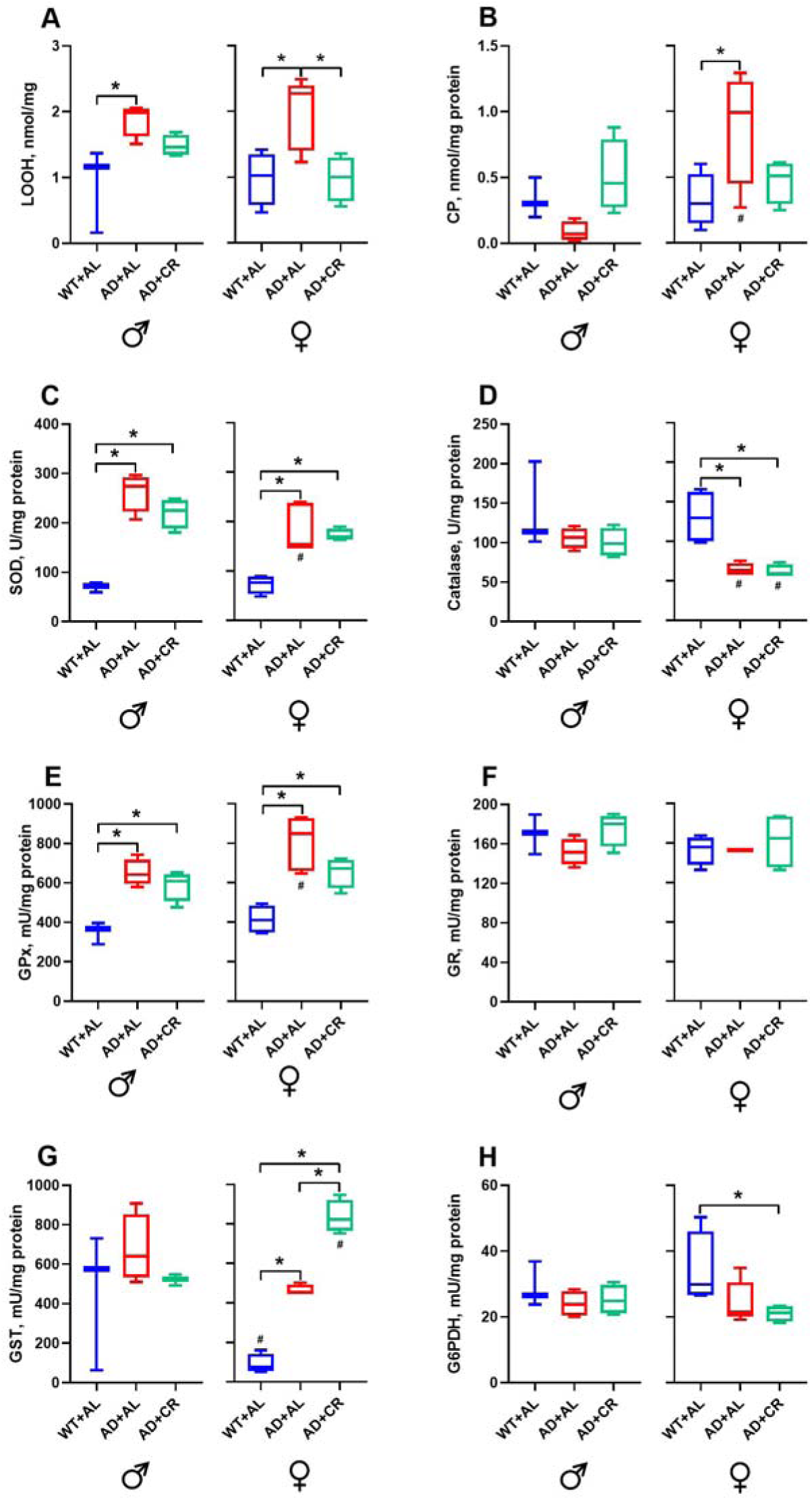
Levels of the oxidative stress markers and activities of antioxidant enzymes in the kidney. A – levels of lipid peroxides, B – levels of carbonyl proteins, C – superoxide dismutase activity, D – catalase activity, E – glutathione peroxidase activity, F – glutathione reductase activity, G – glutathione S-transferase activity, and H – glucose-6-phosphate dehydrogenase activity. *Significantly different between genotype+diet groups, *p <* 0.05. ^#^Significantly different among sexes, *p* < 0.05.

The AL and CR groups of AD males and females had significantly higher kidney SOD activity, compared to the respective WT groups (Fig. 6C). Still, the AD-mediated SOD activity in males was higher than that in females, resulting in a significant sex-specific difference. Compared to the respective WT groups, the catalase activity did not differ in males but was significantly lower in AL (51%) and CR (54%) AD females, again revealing significant sex-specific differences. Compared to respective WT controls, GPx activity was significantly higher in AD+AL and AD+CR groups of both male and female mice (Fig. 6E). Still, GPx activity in AL females was significantly stronger than that observed in AL males. Irrespective of dietary regimen, male AD mice showed no significant differences in kidney GR (Fig. 6F), GST (Fig. 6G), or G6PDH (Fig. 6H) activities compared to WT mice. Similarly, AD females did not differ from WT mice in the activity of kidney GR (Fig. 6F). Surprisingly, in the WT mice, the GST activity was significantly lower (87%) in females compared to males (Fig. 6G). Compared to this control, the GST activity in the kidneys of AD females was 6-fold higher in the AL and 10.9-fold in the CR group (Fig. 6G). Moreover, GST activity was significantly (81%) higher in AD females on the CR regimen compared to the AL group (Fig. 6G). For G6PDH, the only significant difference observed in females was between WT and AD+CR groups (Fig. 6H).

As in the liver (Fig. 5A), glucose levels were significantly higher in the kidneys of male and female AD compared to the respective WT mice (Fig. 7A). AL-fed male AD mice had significantly (3-fold) higher kidney glucose levels than their female counterparts. Moreover, while CR caused a significant (46%) reduction (normalization) of glucose levels in males, it caused a significant (2.2-fold) further increase of glucose levels in females (Fig. 7A).

**Figure 7.**
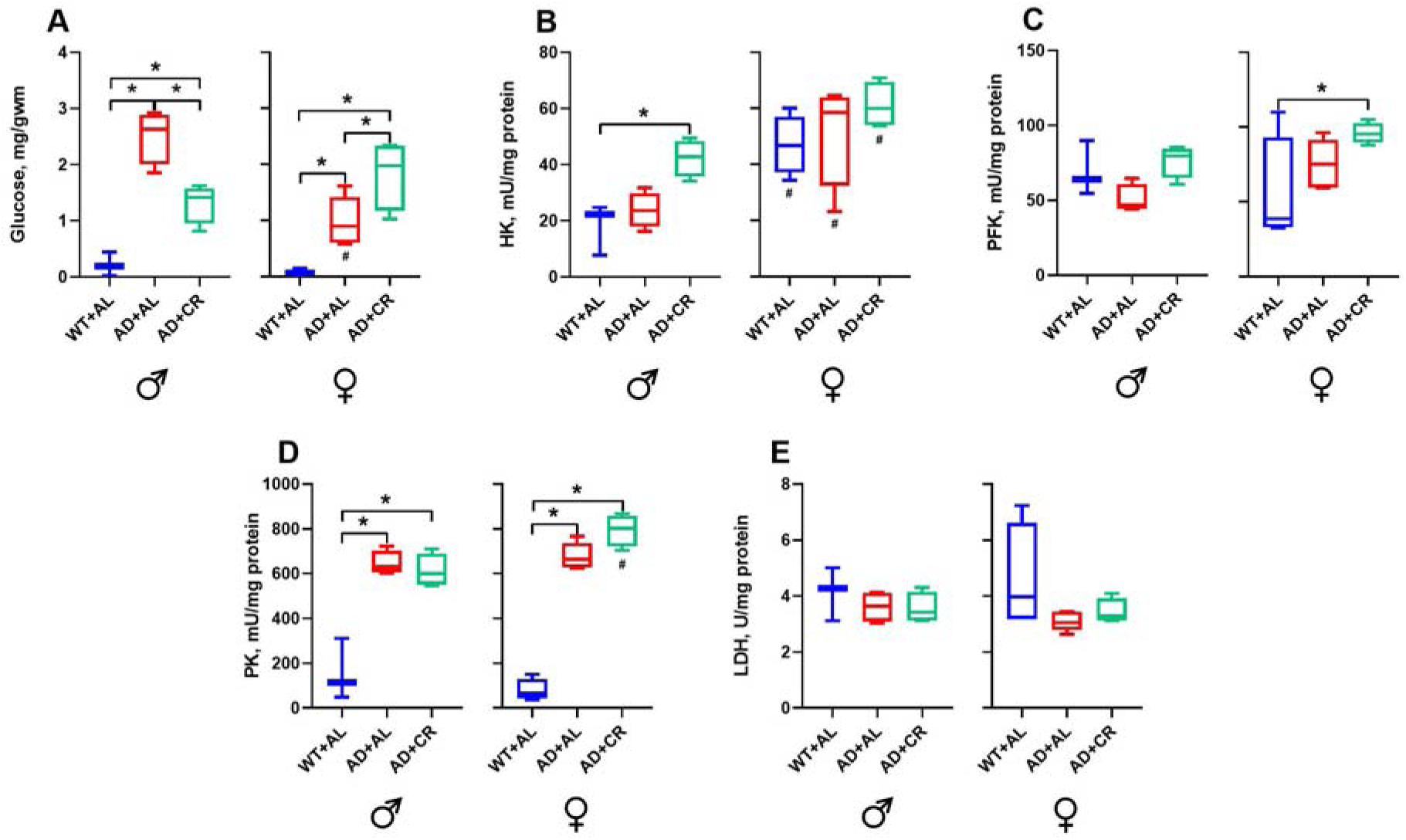
Levels of glucose and activity of glycolytic enzymes in the kidney. A – levels of glucose, B – hexokinase activity, C – phosphofructokinase activity, D – pyruvate kinase activity, and E – lactate dehydrogenase activity. *Significantly different between genotype+diet groups, *p <* 0.05. ^#^Significantly different among sexes, *p* < 0.05.

AL-fed AD males and females had similar to WT mice levels of HK, PFK and LDH activities (Fig. 7B, C, E). Surprisingly, however, HK activity was significantly higher in females compared to males for all groups tested, and the CR regimen selectively increased HK activity in males (Fig. 7B). Compared to WT mice, a small but significant CR-induced increase in PFK activity was observed in AD females (Fig. 7C), while LDH activity remained stable throughout the experiment in both sexes (Fig. 7E). Finally, the kidney of AD males had significantly higher PK activity in AL (5.6-fold) and CR (5.3-fold) groups compared to the WT group (Fig. 7D). Similar effects were also observed in the kidneys of female AD mice, with small but significant sex-specific difference in the AD+CR group.

### 3.4 Principal component analysis

Principal component analyses revealed substantial differences between WT and AD mice in all three organs tested, with male and female data mapping to the overlapping clusters (Fig. 8). In the liver and brain, the effect of caloric restriction was rather subtle. In all organs studied, parameters explaining the largest proportion of variance along PC1 were LOOH, SOD, G6PDH, catalase, and PK. The contribution of several parameters was tissue-specific. Thus, in the brain, along with LOOH, SOD and PK, AD development was associated with large changes in the activity of PFK and HK. There was also a strong association of amyloidosis with the tissue glucose levels, where measured. In the liver, high PFK activity was associated with the WT (Fig. 5C, 8B). In the kidney, GPx and GST, enzymes able to detoxify lipid peroxides, also contributed to the PC1 variance, whereas in the brain and liver, these two enzymes did not show strong associations with either WT or AD phenotype. In the brain and liver, LDH activity explains the largest proportion of variance along PC2, likely indicating the particular importance of this enzyme for these two organs.

**Figure 8.**
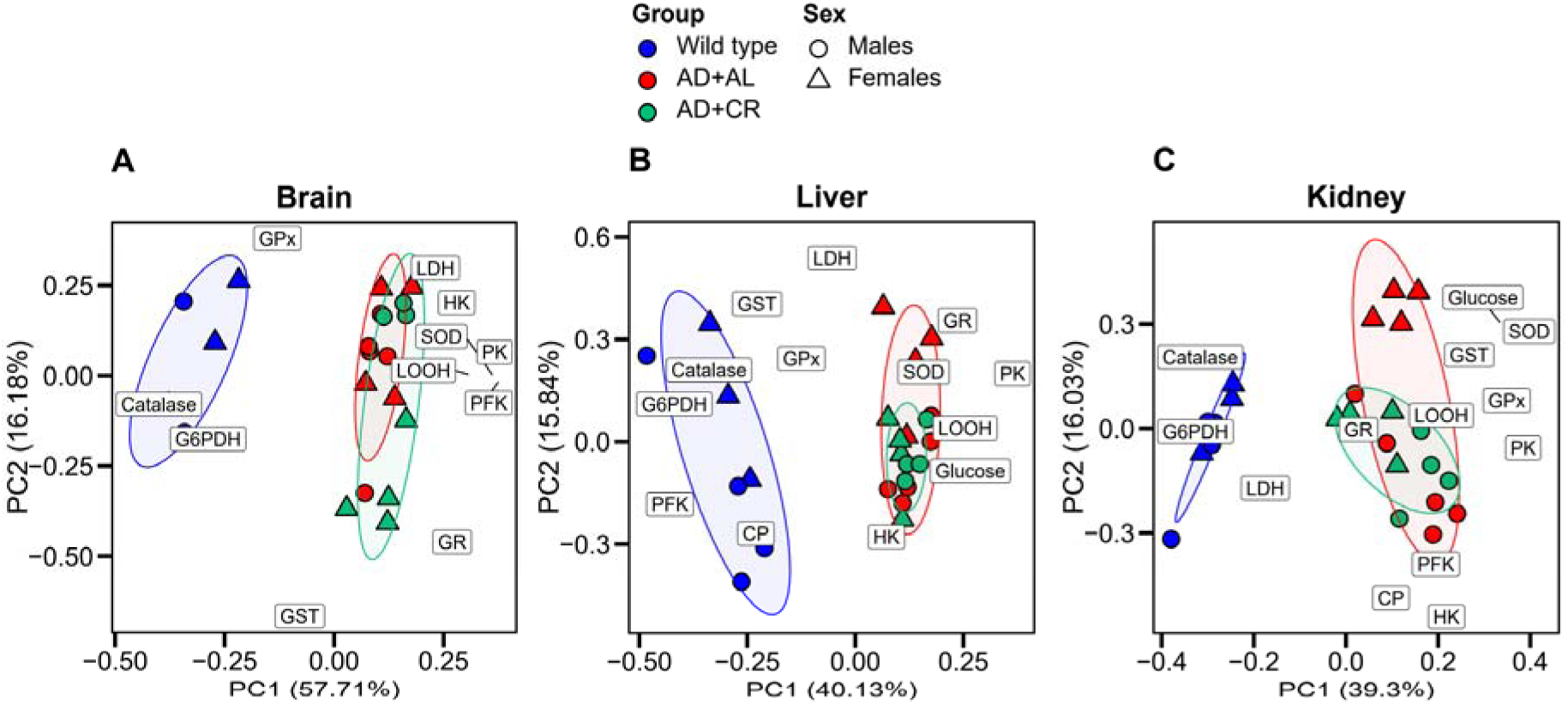
Principal component analysis of all parameters measured in this study. Biplots, demonstrating the contribution of each measured parameter to the total variance of the dataset. The distance of each parameter label from the center of the figure indicates the amount of variance along each principal component. The data used for the PCA analyses were taken from measurements conducted on samples from the brain (A), liver (B), and kidney (C) of AD, AD+CR, and WT mice of both sexes. Note that liver and kidney datasets also comprise measurements of glucose and carbonyl proteins, not present in the brain dataset.

## 4 Discussion

### 4.1 AD-induced body-wide metabolic changes

Recent cumulative evidence refutes the purely central nervous system-centered etiology of AD and positions it as a metabolic disease of the entire body [9,10,39–41]. By analyzing back-to-back AD-induced metabolic changes in the brain, liver, and kidney, i.e., the three key organs for Aβ generation and clearance, we observed a strong and consistent increase in oxidative stress markers (Fig. 2A, 4A, 6A, B). The PCA analysis confirmed a strong association of LOOH levels in all organs studied with the AD phenotype (Fig. 8). The increase in LOOH levels is consistent with previous brain data from humans and different rodent models of AD [5,42]. Thus, neurotoxic byproducts of lipid peroxidation 4-HNE (Fig. 1) and acrolein, were already elevated in the hippocampus, cortex, and cerebellum of subjects with Mild Cognitive Impairment and early AD [43,44]; while high levels of thiobarbiturate-reactive substances and F4-isprostanes were found in the postmortem brains of AD patients [45,46]. Consistently, cortical data obtained in 3-5-month-old female 3×Tg-AD mice suggest that oxidative stress is intensified early during disease progression, before the development of senile plaques or tau pathology, and is likely triggered by the intracellular accumulation of Aβ oligomers [47].

Intracellular Aβ can induce oxidative stress in several ways. In brain mitochondria of AD patients and the mouse model of AD (mAPP mice), it complexes with the Aβ-binding alcohol dehydrogenase, which binds Aβ at nanomolar concentrations [48]. This causes mitochondrial dysfunction, excessive generation of ROS, and cytochrome c release from mitochondria. In addition, Aβ can bind transition metals in a way that favors the production of hydroxyl radicals through the Fenton reaction [49]. Finally, Aβ-induced ROS may interfere with the brain’s antioxidant or glycolytic enzymes, making them more vulnerable to oxidative damage [5]. As the disease progresses, extracellular Aβ oligomers bind to pattern recognition receptors on the immune cells (e.g., Toll-like-receptor 2 and receptor for advanced glycation end-products) on brain microglia [6,50], thus causing the activation of NF-kB, enhanced expression of iNOS (inducible NO-synthase) and NOX2 (NADPH oxidase 2) and increased production of nitric oxide (^•^NO), superoxide anion radical (O_2_^•−^), hydrogen peroxide (H_2_O_2_) and peroxynitrite (ONOO^–^). Moreover, the activation of the NF-kB pathway in immune cells also generates proinflammatory cytokines like interleukin 1β (IL-1β), which, in turn, further activate NF-kB (see Fig. 1 in [6]). Furthermore, extracellular Aβ oligomers induce neuronal hyperactivity and enhance intracellular Ca^2+^ signaling [51]. This, in turn, increases Ca^2+^ content within mitochondrial Ca^2+^ stores [52], whose oxidative phosphorylation (OXPHOS) system and dehydrogenase complexes generate ROS [53]. When intramitochondrial Ca^2+^ levels are high, the Ca^2+^-regulated enzymes α-ketoglutarate dehydrogenase and pyruvate dehydrogenase generate ROS at higher rates [53]. Importantly, in mouse models of AD, and likely also in AD patients, amyloid accumulations are not restricted to the brain but are also found in the liver and kidney [10,54,55].

The three organs under study reacted to the above challenges with an activation of the antioxidant defense, but the activation patterns of antioxidant enzymes turned out to be organ-specific: the activity of (i) SOD was higher in all tissues except the female liver; (ii) catalase was lower in the female brain and kidney as well as in the male liver; (iii) GPx was lower in the female brain and higher in the kidney of both sexes with no change in the liver; and (iv) GST selectively increased in the female kidney.

The AD-mediated decrease in catalase activity might be caused by a possible depletion of active heme, the prosthetic group of catalases, as intracellular Aβ was shown to build complexes with free heme, thereby decreasing its bioavailability [56]. In the brain, liver, and kidney of both sexes, the AD-induced changes in G6PDH activity resembled changes in catalase activity. A strong correlation between the activities of catalase and G6PDH was also found in our previous studies in different model organisms [26,32,57], as G6PDH was shown to be vulnerable to oxidation by ROS, especially when catalase activity is decreased [32,58]. Thus, the ubiquitous AD-mediated decrease in G6PDH activity, visualized also by the PCA biplot (Fig. 8), may represent an additional marker of oxidative stress, along with the increase in LOOH levels and the activity of SOD (as a possible compensatory response). G6PDH is at the crossroads of antioxidant defense (as an enzyme, which produces NADPH, an important reductant) and carbohydrate metabolism. As its activity is lowered in the liver of both sexes and all female tissues studied, carbohydrate metabolism is predominantly directed to glycolysis.

Higher activities of glycolytic enzymes likely reflect the amyloidosis-driven inflammation and metabolic reprogramming of immune cells, shifting from oxidative phosphorylation to glycolysis, even in the presence of oxygen. In peripheral and central immune cells, glycolysis and its metabolites (i) activate proinflammatory signaling pathways promoting, for example, the activation of NFAT (nuclear factor of activated T-cells), NF-κB and mTOR complex 1 (mTORC1) or production of IL-1β and interferon γ; (ii) increase intracellular Ca^2+^ levels and trigger post-transcriptional and post-translational modifications as well as epigenetic changes [59]. In our study, the strongest effect was observed in the brain, with a selective increase in the activities of HK and PFK, accompanied by a ubiquitous increase in the activity of PK. An increased abundance of pyruvate kinase M1 was found previously in cortical extracts of 6-month-old APPswe/PSEN1dE9 male mice [60]. A similar increase was seen in samples of the brain and cerebrospinal fluid of AD patients [61–63]. As glycolytic enzymes are abundantly expressed in the postsynaptic density [64], the higher activity of these enzymes might reflect the ubiquitous AD-induced hyperactivity of neural networks [65,66]. Moreover, microglia are known to undergo Aβ-induced metabolic reprogramming, shifting from OXPHOS to glycolysis to support energy-intensive processes like proliferation, phagocytosis, and production of pro-inflammatory cytokines [67].

PCA analysis indicates an opposite trend for changes in PFK activity in the liver, with the lowest PFK activity found in AD animals (Fig. 5C, 8B). This difference may reflect the predominance of gluconeogenesis in the liver and implies that both gluconeogenesis in the liver and glycolysis in the brain and kidney may help maximize the flux of specific metabolites (e.g., glycerol 3-phosphate). In glycolytic tissues, PFK directs the flow to the production of trioses, whereas in the liver, trioses can be formed in gluconeogenesis, in reverse reactions of the glycolytic pay-off phase. Among trioses, dihydroxyacetone phosphate can be converted into glycerol 3-phosphate, which, in turn, may contribute to the production of ATP by mitochondria through glycerol 3-phosphate shuttle, bypassing complex I of the electron-transport system. This shuttle may provide energy for brain cells [68] and was recently shown to be important for neuronal metabolic flexibility [69].

Collectively, these data show that AD induces a strong and significant metabolic change in the entire body (see also Fig. 8), accompanied by a strong peripheral (and likely also central [70]) hyperglycemia, oxidative damage, as well as the enhancement of glycolysis and antioxidant defense.

### 4.2 Sex-specificity of the AD-induced changes

Women make up almost two-thirds of AD cases, and at the same age (e.g., 65 years of life), women are twice as likely to develop AD as men (<u>Women and Alzheimer’s | Alzheimer’s</u> <u>Association</u>). While the reasons also include longevity, pay gap, and societal and financial stress (e.g., caused by lengthy unpaid caregiving), sex-specific biological differences, including both genetic and sex hormone-driven mechanisms, are crucial and are mimicked by mouse models of the disease [3,71,72]. From the metabolic point of view, the sex-specificity of lipid and glucose metabolism, often connected to neuroinflammation and ROS production, has to be noted [28]. Our data revealed some sex-specific differences already in (middle-aged) WT mice. Thus, the activity of catalase differed between males and females, both in the brain and the liver, being, however, higher in the female brain and the male liver. Besides, LDH activity was higher in the male liver, and that of HK - in the female kidney. All but one (kidney HK activity) of the mentioned above differences disappeared in AD mice, showing that similarities between sexes under disease conditions can have different etiologies. Instead, compared to male AL-fed AD mice, PK and PFK activities were lower in the brains of females, accompanied by a lower SOD and higher HK activities in the liver and kidney. Besides, the G6PDH activity was consistently lower in AD than in WT females, resulting in decreased NADPH production. In AD mice, kidneys turned out to be the most sex-sensitive organ, having also lower glucose and higher protein carbonyl levels as well as lower catalase activities in females compared to males.

Together, these data point to less efficient AD-induced glycolysis in the female brain (and, consequently, reduced inflammatory response [59]), as well as lower efficiency of antioxidant defense in the female liver and kidney (likely impacting the Aβ clearance), thus providing yet another facet to the complex picture of female vulnerability in AD.

### 4.3 Impact of caloric restriction

Obesity, insulin resistance, and type 2 diabetes are well-known comorbidities of AD sharing common cellular/molecular markers like oxidative damage, neuroinflammation, Ca^2+^ dyshomeostasis, dysregulation of neural network activity, and impaired autophagy, adult neurogenesis, as well as learning and memory [73]. At the same time, caloric restriction prolongs the lifespan of different species from yeast to humans; upregulates autophagy and DNA repair; suppresses oxidative stress and neuroinflammation; stabilizes neuro-glial Ca^2+^ homeostasis and neural network activity; stimulates mitochondrial biogenesis and neurogenesis [73,74]. Moreover, CR was shown to decrease amyloid deposition in mouse models of AD [16,18] and improve the performance of elderly subjects in the auditory verbal learning task [19]. Therefore, we expected CR to have a profound effect, counteracting the AD-induced metabolic dysfunction. To our surprise, the levels of oxidative stress markers were reduced by CR only in the female kidneys. In the brain, LOOH levels remained unchanged, whereas in the liver, they were significantly higher compared to those found in AL-fed AD mice. In the brain, however, CR sex-specifically increased the activity of antioxidant enzymes, enhancing the activity of SOD and GPx in females as well as GR and to a lesser extent GST in males. In the liver, CR brought the levels of male SOD as well as female GPx and GR, close to those observed in WT mice. The only CR effect observed in the kidneys was a further increase in the activity of female GST compared to AL-fed AD mice.

For glycolytic enzymes, we noticed a CR-induced increase in female PFK activity and a decrease in male PK activity in the brain, as well as an increase in female PK activity in the liver. Under the caloric restriction, AD females had significantly higher PFK activity in the brain, HK activity in the liver and kidneys, as well as PK activity in the kidneys, compared to their male counterparts. Thus, in females, glycolysis is more responsive to CR than in males, so that CR-induced enhancement of glycolysis might help to regulate AD-induced inflammation [59]. In general, however, all observed differences between the AL-fed and calorie-restricted AD mice were much smaller than the ones between WT and AD mice (Fig. 8).

## 5 Conclusions

The current study shows that AD-related mutations in genes encoding amyloid precursor protein and presenilin 1 induce body-wide adverse changes, increasing the levels of lipid peroxides in the brain, liver, and kidney, and strongly affecting the activities of the first-line antioxidant enzymes, such as superoxide dismutase and catalase. The changes in the activities of these enzymes are not unidirectional: mutant mice exhibit higher superoxide dismutase, but lower catalase activity compared to their WT counterparts. In all tissues studied, AD phenotype was associated with higher activity of pyruvate kinase, whereas differences in activities of glutathione-related enzymes, and enzymes of glucose catabolism, glucose 6-phosphate dehydrogenase, hexokinase, and phosphofructokinase, between the WT and mutant mice were tissue- and sex-specific. Under caloric restriction, female mice had more effective glucose catabolism, and their kidneys showed the strongest decrease in the oxidative stress markers among all studied tissues. In general, however, the impact of caloric restriction on key glycolytic, antioxidant, and related enzymes was rather modest.

## Acknowledgements

We thank E. Zirdum and K. Schmidt for technical assistance, and E. Zirdum, K. Pan and A. Hahn for the help with caloric restriction and tissue sampling.

## CRediT authorship contribution statement

**Myroslava V. Vatashchuk:** investigation, visualization, writing – original draft (Sections “Methods and Materials”, “Results”)

**Viktoriia V. Hurza:** investigation

**Maria M. Bayliak:** validation, data curation, writing – review and editing

**Dmytro V. Gospodaryov:** formal analysis, writing – original draft, writing - review and editing

**Volodymyr I. Lushchak:** conceptualization; methodology; writing – review and editing

**Olga Garaschuk:** conceptualization; methodology; supervision; writing – review and editing; funding acquisition.

## Data availability

Data will be made available on request.

## Abbreviations

AD: Alzheimer’s disease
WT: wild-type
AL: ad libitum
?R: caloric restriction
APP: amyloid precursor protein
PS1: presenilin 1
A?: amyloidROS reactive oxygen species
LOOH: lipid peroxides
CP: carbonyl proteins
SOD: superoxide dismutase
G6PDH: glucose-6-phosphate dehydrogenase
GPx: glutathione peroxidase
GR: glutathione reductase
GST: glutathione S-transferase
HK: hexokinase
LDH: lactate dehydrogenase
PFK: phosphofructokinase
PK: pyruvate kinase

